# Epigenomic analysis of Parkinson’s disease neurons identifies *Tet2* loss as neuroprotective

**DOI:** 10.1101/779785

**Authors:** Marshall Lee, Killinger Bryan, Li Peipei, Ensink Elizabeth, Li Katie, Cui Wei, Lubben Noah, Weiland Matthew, Gordevicius Juozas, Coetzee Gerhard A., Jovinge Stefan, Labrie Viviane

## Abstract

PD pathogenesis may involve the epigenetic control of enhancers that modify neuronal functions. Here, we comprehensively profile DNA methylation at enhancers, genome-wide, in neurons of 57 PD patients and 48 control individuals. We found a widespread increase in cytosine modifications at enhancers in PD neurons, which is partly explained by elevated hydroxymethylation levels. Epigenetic dysregulation of enhancers in PD converge on transcriptional abnormalities affecting neuronal signaling and immune activation pathways. In particular, PD patients exhibit an epigenetic and transcriptional upregulation of *TET2*, a master-regulator of cytosine modification status. *TET2* inactivation in a neuronal cell line results in cytosine modification changes that are reciprocal to those observed in PD neurons. Furthermore, *Tet2* inactivation in mice fully prevents dopaminergic neuronal loss in the substantia nigra induced by prior inflammation. *Tet2* loss in mice also attenuates transcriptional immune responses to an inflammatory trigger. Thus, widespread epigenetic dysregulation of enhancers in PD neurons may, in part, be mediated by increased *TET2* expression. Decreased *Tet2* activity is neuroprotective, *in vivo*, and may be a novel therapeutic target for PD.

## Introduction

Parkinson’s disease (PD) is a severely debilitating neurodegenerative disorder involving progressive motor and non-motor symptoms that affects more than six million individuals worldwide^1^. The vast majority of PD cases are idiopathic with unclear etiopathological mechanisms, though this disease has numerous clinical manifestations that are consistent with an epigenetic basis. Epigenetic mechanisms are a partially dynamic means of gene regulation that represent a convergence point for environmental, aging and genetic risk factors. Clinical features of PD such as the delayed age of disease onset, low concordance rate (11%) between monozygotic twins^2, 3^, symptomatic fluctuations over disease course, including a unilateral to bilateral symptom presentation in most individuals (85%)^4, 5^, and prominent sex differences in disease risk^6, 7^, collectively suggest the contribution of the modifiable epigenome to disease origins and progression^8–10^. Although recent candidate-gene and array-based studies support that epigenetic abnormalities occur in PD tissues^10–14^, thus far, the epigenomes of PD patients remain largely unexplored and there has yet to be an epigenetic study of neurons from the PD brain.

Early synaptic dysfunction, neuronal deposits of aggregated α-synuclein, and gradual neurodegeneration of the nigrostriatal system are all hallmark pathological features of PD that evince the involvement of neurons in this disease^15^. The epigenetic modification DNA methylation plays a critical role in activity-dependent neuronal gene expression and neuronal survival^16, 17^. DNA methylation also undergoes widespread remodeling with age^18–20^ and in response to environmental toxins associated with PD^21, 22^. The conversion of DNA methylation to hydroxymethylation (and eventual demethylation) is catalyzed by ten-eleven translocation (TET) enzymes, which have also been shown to impact neurodevelopment and synaptic transmission^18, 23–25^. Despite their roles in healthy neuronal functions, there have been relatively few genome-wide studies of DNA methylation in the PD brain^11–13^. These have focused on bulk brain tissue with a heterogeneous mixture of neurons and glial cells, using platforms that examine primarily CpG sites within coding regions (exons) and CpG islands^11–13^. However, it is now established that DNA modification profiles strongly diverge between neurons and glial cell types, and a large proportion of DNA methylation in neurons occurs at non-CpG (CpH) sites^18, 26, 27^. Furthermore, there are numerous important regulatory elements of genes, such as enhancers, located outside of the regions previously examined that may play an important role in PD^28^.

Enhancers are cell-type specific elements that control complex gene expression patterns, and their activation state is regulated by epigenetic marks^28, 29^. Genetic studies point to disrupted enhancer function in PD, affecting dopaminergic neurons^30, 31^ and the regulation of PD genes, including α-synuclein^32, 33^. DNA methylation analysis of dopamine neurons derived from induced pluripotent stem cells from PD patients also find a strong enrichment of enhancer disruption in PD^34^. Thus, an in-depth analysis of epigenetic regulation at enhancers genome-wide in mature neurons isolated from the PD brain has considerable potential for uncovering pathobiological processes in PD.

Here, we fine-mapped DNA methylation and hydroxymethylation at enhancers genome-wide in neurons of the prefrontal cortex of PD patients and controls (n=57 and 48 individuals, respectively). Remarkably, enhancers in PD neurons exhibit a widespread increase in cytosine modifications that is, in part, the result of elevated levels of hydroxymethylation. Chromatin conformation analysis reveals that genes with an enhancer disruption are involved in neurogenesis. Transcriptome-wide analysis supports the dysregulation of genes with epigenetically altered enhancers in PD, and further demonstrates an activation of neurodevelopmental and immune pathways in the PD brain. Notably, there is an epigenetic misregulation and transcript upregulation of the *TET2* gene in PD. Enhancer-wide hypermethylation in PD neurons is linked to *TET2* disruption, as shown in an *in vitro* model. Moreover, we find that *Tet2* inactivation has neuroprotective effects on nigral dopaminergic neurons in aged mice previously exposed to inflammation. Together, our study suggests a widespread epigenetic disruption of enhancers in PD neurons that is partially mediated by increased *TET2*, and that silencing *TET2* in PD patients may be a novel neuroprotective therapy.

## Results

### Epigenetic disruption of enhancers in PD neurons

To identify epigenetically misregulated enhancers involved in PD, we performed a genome-wide analysis of DNA methylation at enhancers comparing neurons of 57 PD patients to those of 48 controls (Supplementary Data 1). Neuronal nuclei were isolated from the prefrontal cortex using fluorescence-activated cell sorting (Supplementary Fig. 1). We then used a targeted bisulfite deep sequencing approach, known as bisulfite padlock probe sequencing, involving 59,064 probes to profile 31,590 brain enhancers, genome-wide. The brain enhancers profiled were defined by the NIH Epigenomics Roadmap, and included all active, poised/bivalent, weak, and genic enhancers, as well as promoters since these have the potential to act as enhancers^29^ (ChromHMM 18-states model). In total, we profiled 904,511 modified cytosine sites at enhancers, examining both CpG and CpH sites (111,412 CpG and 793,099 CpH sites assessed; Supplementary Fig. 2). The prefrontal cortex is composed of glutamatergic and GABAergic neurons, and we performed a neuronal subtype deconvolution with reference markers^35, 36^ to confirm that there was not a selective loss of neuronal subtypes between PD patients and controls (Supplementary Fig. 3). We also adjusted for the proportion of glutamate to GABAergic neuronal levels in our analysis. Differentially methylated cytosines in PD neurons were identified using a logistic regression model, adjusting for age, sex, postmortem interval, and neuronal subtype proportion.

There were 1,799 differentially methylated cytosines at enhancers in PD neurons compared to healthy control neurons (*q*<0.05, logistic regression; **Fig. 1a**; Supplementary Data 2). The vast majority of enhancers in PD neurons were hypermethylated (74.3%; 1,337 out of 1,799 cytosines; OR=2.3, *p*<10^-15^, Fisher’s exact test; **Fig. 1b**). Most of the differentially methylated cytosines in PD neurons were distally located from transcription start sites (average distance: 48.49 kb ± 0.169 s.e.m.; Supplementary Fig. 2c), supporting that enhancers were predominantly epigenetically altered in PD neurons. Of the differentially methylated sites, there were 284 at CpG and 1,515 at CpH sites (average cytosine modification change: 7.33% ± 0.171 s.e.m and 1.77% ± 0.057 s.e.m at CpG and CpH sites, respectively), with CpA sites being most frequently disrupted at enhancers in PD (**Fig. 1c**). Transcription factor motif analysis of the differentially methylated sites revealed an enrichment of the Ras-responsive binding protein 1 (RREB1; **Fig. 1d**), which is involved in axonal degeneration^37^ and enables the activity of PD-relevant genes like Parkinsonism-associated deglycase (*DJ-1*)^38^.

**Fig. 1.**
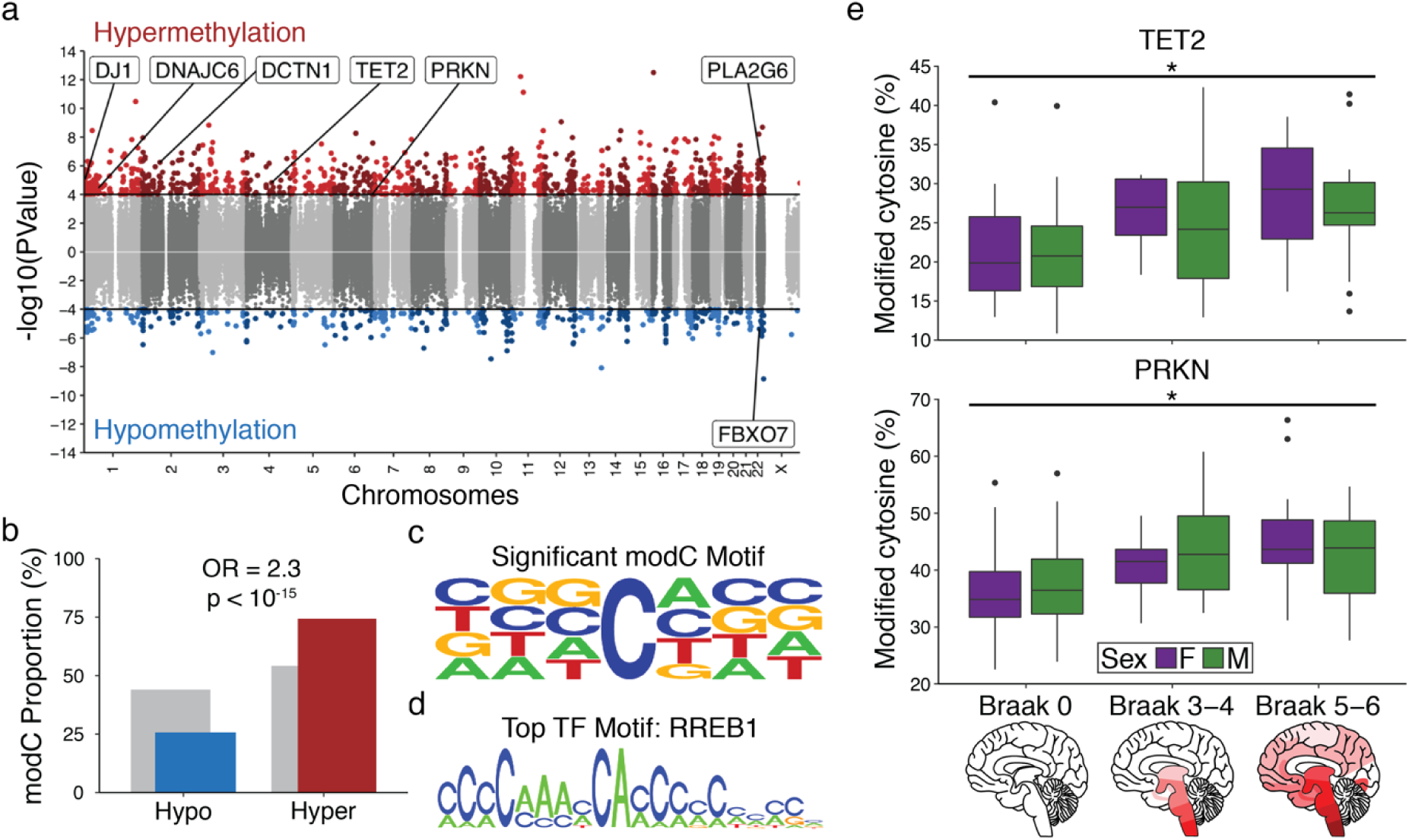
Genome-wide hypermethylation at enhancers in PD prefrontal cortex neurons. (**a**) Manhattan plot showing differentially methylated cytosines at enhancers in PD, identified by logistic regression after adjusting for age, sex, postmortem interval, and neuronal subtypes (n=57 PD, 48 controls). -log10(PValue) refers to the significance of cytosine methylation change in PD, with the sign corresponding to the direction of change (red: hypermethylated, blue: hypomethylated cytosine in PD). Threshold for genome-wide significance is *q*<0.05 (black line). Differentially modified cytosines at enhancers associated with known familial PD genes are highlighted. (**b**) Proportion of hypermethylated (red) and hypomethylated (blue) cytosines at enhancers in PD. Enrichment determined by Fisher’s exact test with OR referring to the odds ratio and *p*<10^-15^ is significance of hypermethylation enrichment. Gray is all cytosines in the analysis (background). (**c**) Motif analysis using Seqlogo depicting contribution of CpG and CpH sites to epigenetic changes in PD, with CpA being the major contributor. (**d**) Top transcription factor motif associated with differentially methylated cytosines in PD, as determined by oPOSSUM. (**e**) Boxplots of the most differentially methylated cytosine in *TET2* and *PRKN* across PD Braak Stages (n=48 controls (Braak stage 0), 22 moderate PD (Braak stage 3-4), 34 advanced PD (Braak stage 5-6)). \**q*<0.05 determined by logistic regression after adjusting for age, sex, postmortem interval, and neuronal subtypes. Boxplot center line is the mean, the lower and upper limits are the first and third quartiles (25th and 75th percentiles), and the whiskers are 1.5× the interquartile range.

We next determined the extent to which enhancers are disrupted early in PD, prior to the arrival of the hallmark Lewy pathology that contains aggregated α-synuclein. In PD, motor symptoms become apparent at Braak stage 3-4, when Lewy pathology enters the substantia nigra and there is nigral dopaminergic neuron loss, but pathology has not yet reached the neocortex^15^. In advanced PD, Braak stage 5-6, disease pathology has progressed to the neocortex, including the prefrontal cortex^15^. Comparisons between PD Braak stage 3-4 and controls showed that there were 2,172 differentially methylated cytosines in enhancers in prefrontal cortex neurons prior to the arrival of PD pathology (*q*<0.05, logistic regression; Supplementary Fig. 4; Supplementary Data 3); these changes largely involved hypermethylation (1,910 out of 2,172 sites, OR=1.7, *p*<10^-15^, Fisher’s exact test). Differentially methylated cytosines in early PD (stage 3-4) also strongly overlapped those identified in the entire PD cohort analysis (607 out of 1,799 sites, OR=194, *p*<10^-15^, Fisher’s exact test). There were also 299 DNA methylation differences common between early and late stage PD, indicating continued epigenetic disruption of these enhancers (*q*<0.05, logistic regression; Supplementary Fig. 4; Supplementary Data 3). Together, this supports a prominent dysregulation of enhancers in PD neurons, occurring prior to onset of PD pathology.

### Gene targets of epigenetically altered enhancers in PD

We sought to understand the effects of widespread hypermethylation of enhancers in PD by identifying the gene targets of the differentially methylated enhancers in PD. Enhancers activate genes through chromatin loops that allow physical interaction between enhancers and their target gene promoters^39, 40^. Analysis of 3D chromatin architecture in the prefrontal cortex, using Hi-C data^41^, revealed that the enhancers with differential methylation in PD targeted 2,469 gene promoters (TSS ± 2 kb; **Fig. 2a**; Supplementary Data 4). To further capture proximal enhancer-promoter interactions, we used an *in silico* cis-regulation prediction tool. In total, there were 2,885 genes affected by the differentially methylated enhancers in PD neurons. These genes strongly affect pathways involved in neurogenesis, neurodevelopment, and synaptic structure (*q*<0.05, hypergeometric distribution; **Fig. 2b**). Furthermore, genes with enhancer perturbations were implicated in neurocognitive and neurodegenerative diseases (top 10 disease pathways; *q*<0.05, hypergeometric distribution; **Fig. 2b**).

**Fig. 2.**
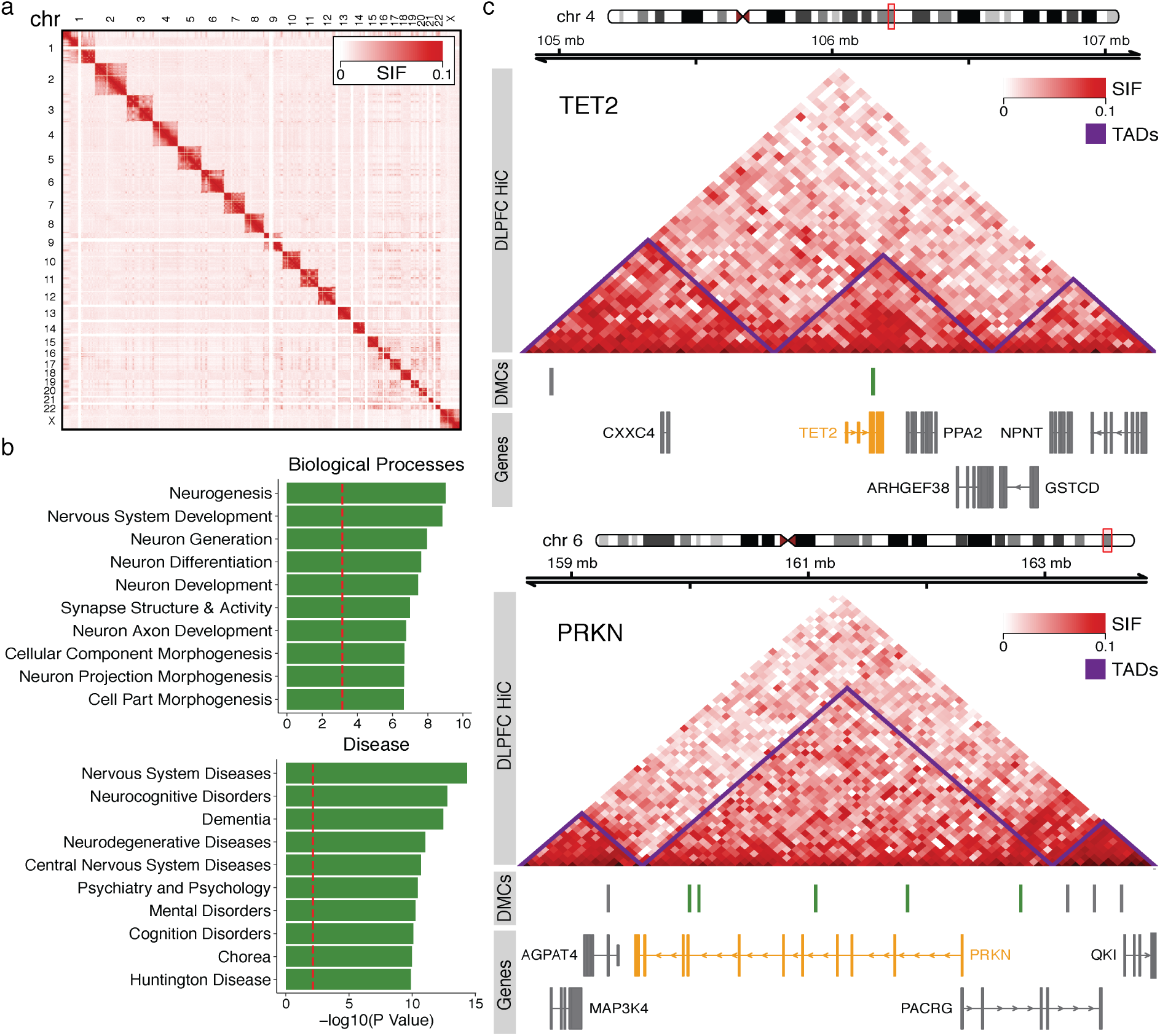
Chromatin conformation analysis identifying gene targets of enhancers that are epigenetically dysregulated in PD neurons. (**a**) Overview of the genome-wide Hi-C interactions in human prefrontal cortex tissue^41^. Heatmap shows interaction frequency with a 500 kb resolution (n=4 individuals). (**b**) Top 10 biological and disease pathways affected by genes with differential methylation at enhancers in PD, as determined by MetaCore. -log10(PValue) refers to the enrichment of pathways in PD, with the red dashed line being *q*<0.05, as determined by hypergeometric test. (**c**) Chromatin conformation analysis revealing enhancer-gene target interactions for *TET2* (upper panel) and *PRKN* (lower panel). In the interaction heat map, red signals a high signal interaction frequency (SIF) and topologically associated domains (TADs) are indicated in purple. Interaction resolution at 40 kb. Differentially methylated cytosines at enhancers in PD neurons in the *TET2* and *PRKN* gene locus (green bars) and surrounding area (gray bars) are shown.

Interestingly, we found an epigenetic disruption of an enhancer targeting the *TET2* gene ‒ a key regulator of DNA methylomes, as it encodes an enzyme responsible for the oxidation of DNA methylation to hydroxymethylation and the removal of DNA methylation^23, 42^ (**Fig. 1e and 2c**). We also found several familial PD risk genes with epigenetically dysregulated enhancers, such as *DJ-1*, auxilin (*DNAJC6*), dynactin 1 (*DCTN1*), parkin (*PRKN*), phospholipase A2 group VI (*PLA2G6*), and F-Box protein 7 (*FBXO7*). Among the risk genes identified for PD by a large meta-analysis of genome-wide association studies (GWAS), 15 genes had a differentially methylated enhancer in PD neurons (out of the profiled 61 PD GWAS-identified genes with SNPs at *p*<10^-8^)^43–45^. Moreover, we found that many of these genes had differential methylation at their enhancers that was related to the progression of Lewy pathology in the PD brain (*q*<0.05, logistic regression; Supplementary Data 4). For example, *TET2* and *PRKN* exhibited DNA hypermethylation at enhancers before and after the arrival of cortical Lewy pathology (**Fig. 1e and 2c**).

### Hydroxymethylation at enhancers in PD neurons

Since we observed *TET2* enhancer dysregulation, we asked whether there were changes in levels of 5-hydroxymethylcytosine in PD neurons. We profiled hydroxymethylation genome-wide in neurons of PD patients and controls (n=21 and 23 individuals, respectively), using hMeDIP-seq. Hydroxymethylated peaks significantly associated with PD were enriched in the gene body, promoters, and enhancers (*p*<0.05, Fisher’s exact test; **Fig. 3a**; Supplementary Data 5). Interestingly, these genomic locations exhibited a significant gain in hydroxymethylation in PD patients (*p*<0.05, Fisher’s exact test; **Fig. 3b**). We next determined the extent to which hyper-hydroxymethylation in PD neurons contributed to the observed enhancer hypermethylation in patients described above. For this, we profiled hydroxymethylation at enhancer cytosine sites associated with PD in our targeted bisulfite sequencing study. There was a strong increase in hydroxymethylation at enhancers in PD neurons (OR=1.95, *p*<10^-15^, Fisher’s exact test; **Fig. 3c**; Supplementary Data 6). Of the enhancers with hydroxymethylation changes in PD neurons (nominal *p*<0.05 and log-fold change≥1, logistic regression, after adjustment for age, sex, postmortem interval, and neuronal subtype proportion), 72.6% exhibited a hyper-hydroxymethylation (OR=1.96, *p*<10^-15^, Fisher’s exact test; **Fig. 3d and 3e**; Supplementary Data 7). Together, this suggests that the widespread DNA hypermethylation observed at enhancers in PD neurons is, in large proportion, the result of a gain in hydroxymethylation, which may be due to the disrupted activity of *TET2*.

**Fig. 3.**
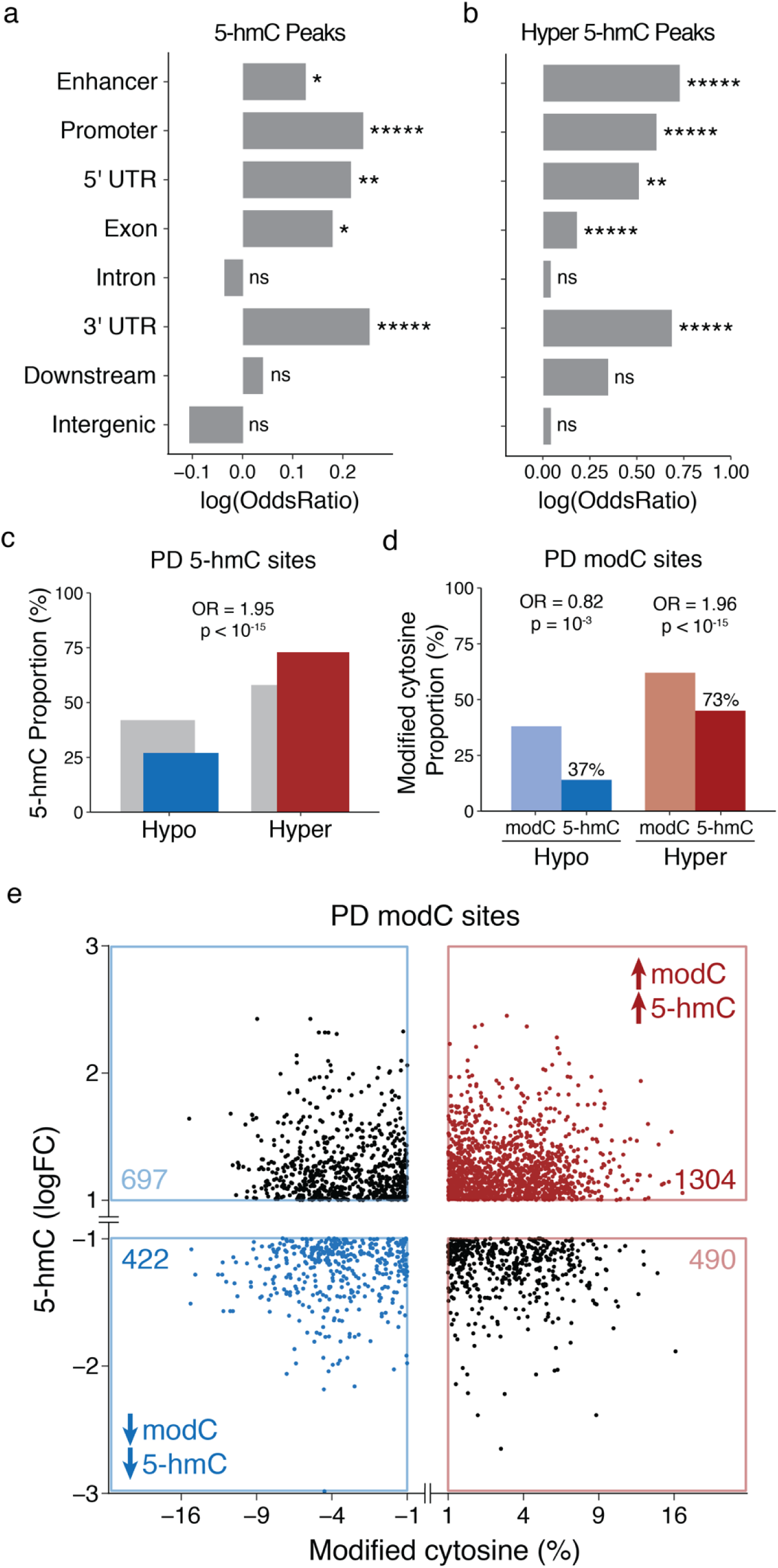
Increased 5-hydroxymethylation contributes to epigenetic abnormalities at enhancers in PD neurons. Differential hydroxymethylation peaks in PD were identified using logistic regression, adjusting for age, sex, postmortem interval, and neuronal subtypes (n=21 PD, 23 controls). (**a, b**) Genomic elements exhibiting hydroxymethylation changes in PD. Genomic elements with an overrepresentation of differential hydroxymethylation (**a**) or hyper-hydroxymethylation (**b**) in PD neurons are shown. Enrichment score reported as log(OddsRatio) and \**p*<0.05, \*\**p*<0.01, \*\*\*\*\**p*<0.00001, as determined by Fisher’s exact test. (**c**) Direction of hydroxymethylation change in PD neurons, genome-wide. Enrichment of hyper-hydroxymethylation in PD genome-wide determined by Fisher’s exact test and shown by odds ratio (OR) and *p*<10^-15^. Gray is all cytosines in analysis (background). (**d**) Contribution of hydroxymethylation to epigenetic changes at enhancers in PD neurons. Percent overlap of hydroxymethylation and direction of hydroxymethylation change is shown for enhancer cytosine sites that were epigenetically altered in PD neurons (nominal *p*<0.05 cytosine sites identified in Fig. 1). Enrichment of hydroxymethylation at epigenetic misregulated enhancers in PD determined by Fisher’s exact test. (**e**) Scatter plot showing the contribution of hyper-hydroxymethylation to epigenetic changes at enhancers in PD neurons. Hydroxymethylation levels at enhancer sites identified to be differentially methylated in PD (nominal *p*<0.05 cytosines in Fig. 1). Hyper-hydroxymethylation occurring at sites with increased cytosine modifications in PD neurons shown in red and hypo-hydroxymethylation occurring at sites with decreased cytosine modifications in PD neurons shown in blue.

### Transcriptional changes in PD

We examined whether the DNA methylation (and hydroxymethylation) changes observed in PD were linked to changes in gene transcript levels. We profiled transcriptomic alterations in the PD prefrontal cortex by RNA-sequencing, using a subset of the same samples from our epigenetic analysis (n=12 controls and 24 PD). Our analysis identified 1,032 differentially expressed genes in PD, after correcting for age, sex, RIN, and sources of unknown variation (*q*<0.05, logistic regression; **Fig. 4a**; Supplementary Data 8). Pathway analysis that integrated transcriptomic alterations with the DNA methylation changes showed prominent immune activation, impaired synaptic signaling, apoptosis-related peptidase activity, and altered neuronal differentiation (*q*<0.01, GSEA; **Fig. 4b**; Supplementary Data 9).

**Fig. 4.**
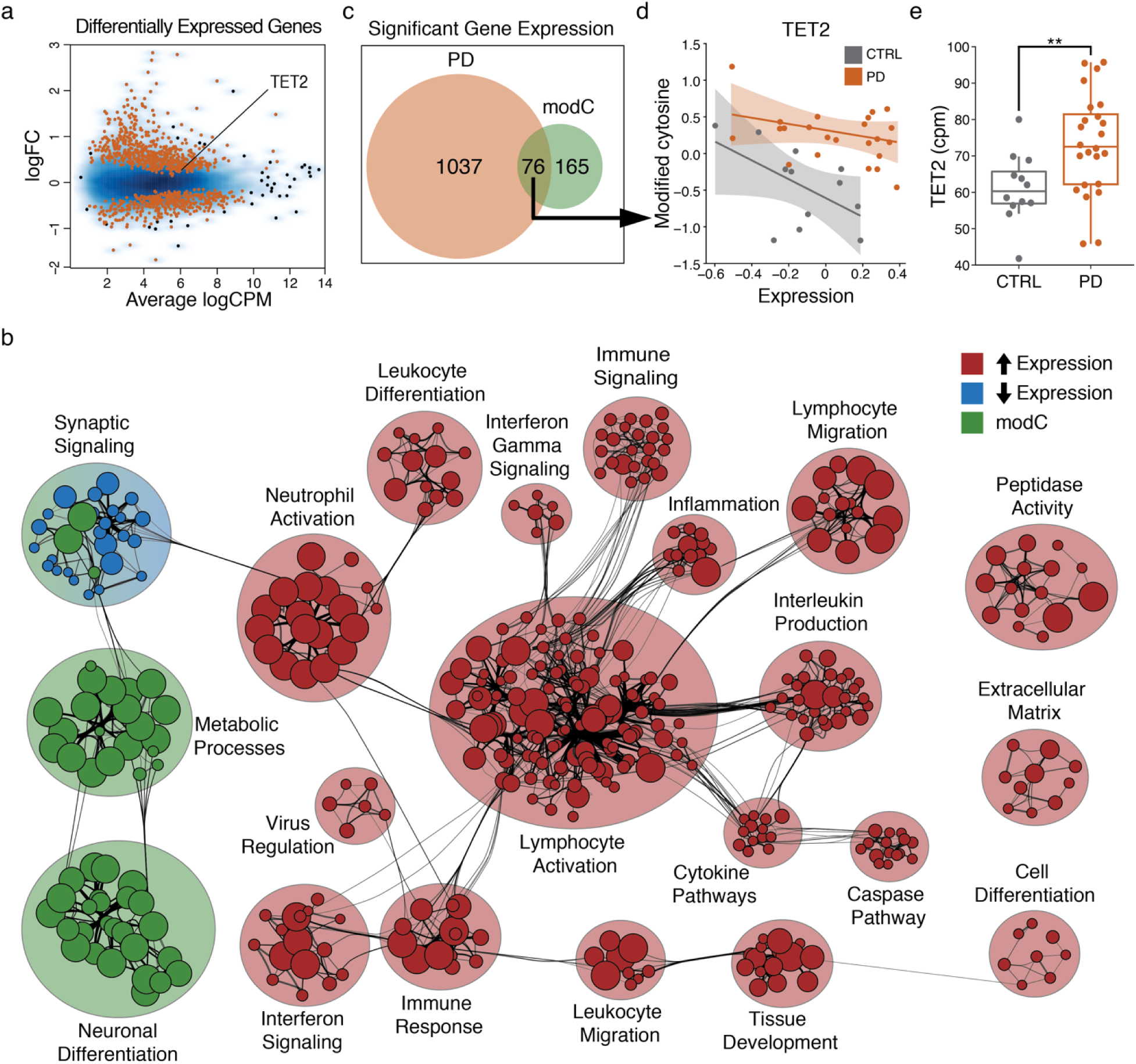
Transcriptomic analysis reveals an association between gene expression and DNA methylation changes in PD that impact immune activation and neuronal signaling. Differentially expressed genes in PD were identified using a generalized linear model, adjusting for age, sex, RIN, and sources of unknown variation (n=24 PD, 12 Controls). RNA-seq data was also compared to DNA methylation data for the same individuals (cytosine methylation at enhancers adjusted for age, sex, postmortem interval, and neuronal subtypes). (**a**) Transcript level changes in the PD prefrontal cortex determined by RNA-seq. Significant changes in gene transcript levels in PD highlighted in red (*q*<0.05, logistic regression). (**b**) Pathway analysis merging transcriptional and DNA methylation changes in PD prefrontal cortex, as determined by GSEA and gProfileR, respectively (*q*<0.01). (**c**) Venn diagram of genes with transcript levels associated with PD and/or enhancer DNA methylation status. *TET2* transcript levels were associated with PD and enhancer DNA methylation (denoted by arrow). (**d**) The relationship between *TET2* gene expression and *TET2* enhancer cytosine methylation in PD patients (gray) and controls (orange). Scatter plot of the correlation between residual *TET2* expression and enhancer DNA methylation status with confidence intervals. (**e**) *TET2* expression in the prefrontal cortex of PD patients compared to healthy controls. Transcript levels in counts per million (cpm) in RNA-seq data. \*\**p*<0.01 after adjusting for age, sex, RIN, and sources of unknown variation. Boxplot center line is the mean, the lower and upper limits are the first and third quartiles (25th and 75th percentiles), and the whiskers are 1.5× the interquartile range.

In PD, genes with epigenetically misregulated enhancers showed a significant enrichment in differential expression (OR=1.20, *p*<0.05, Fisher’s exact test; Supplementary Data 10). Of the 241 genes that showed a significant correlation between their enhancer DNA methylation status and gene transcript levels, there were 76 that were both epigenetically and transcriptionally dysregulated in PD (**Fig. 4c**; Supplementary Data 11). *TET2* was one of these 76 genes, exhibiting a significant inverse correlation between enhancer DNA methylation status and *TET2* transcript levels (**Fig. 4d**). We also found that in the PD prefrontal cortex, *TET2* levels were increased relative to controls (nominal *p*<0.05, logistic regression; **Fig. 4e**). No changes in *TET1* or *TET3* were observed in the PD prefrontal cortex (Supplementary Fig. 5). Thus, elevated levels of *TET2* may underlie the abnormalities in hydroxymethylation/methylation at enhancers in PD neurons.

### Validation and consequences of epigenetic disruption of TET2 in PD

We further confirmed and characterized *TET2* dysregulation in neurons of the PD prefrontal cortex. We first analyzed *TET2* transcript levels in isolated neuronal and glial (non-neuronal) nuclei, as well as in the bulk cytosolic fraction. *TET2* mRNA levels were elevated in PD, particularly in neurons and in the bulk cytosol, relative to controls (main effect of diagnosis: F(1,36)=2.60, *p*<0.05, repeated-measures ANOVA; n=10 PD and 10 controls; **Fig. 5a**). Next, to determine the full extent of epigenetic perturbations at the *TET2* gene in prefrontal cortex neurons of PD patients, we mapped DNA methylation at the entire *TET2* gene locus and surrounding genomic area (± 300 kb) in PD patients (n=20) and controls (n=11) using the targeted bisulfite approach. There were 15 differentially methylated cytosine sites in the *TET2* region in PD neurons, after adjusting for age, sex, postmortem interval, and neuronal subtype proportion (*q*<0.05, logistic regression; **Fig. 5b**; Supplementary Data 12). Notably, we found pronounced hypermethylation of an enhancer in the *TET2* gene body; however the promoter of *TET2* was hypomethylated (*q*<0.05, logistic regression; **Fig. 5b**). Furthermore, we observed that DNA methylation alterations in *TET2* increased with the severity of PD pathology (*q*<0.001, logistic regression; 15.9% methylation difference; **Fig. 1e**), supporting the dysregulation of this epigenome-modifying enzyme in PD neurons.

**Fig. 5.**
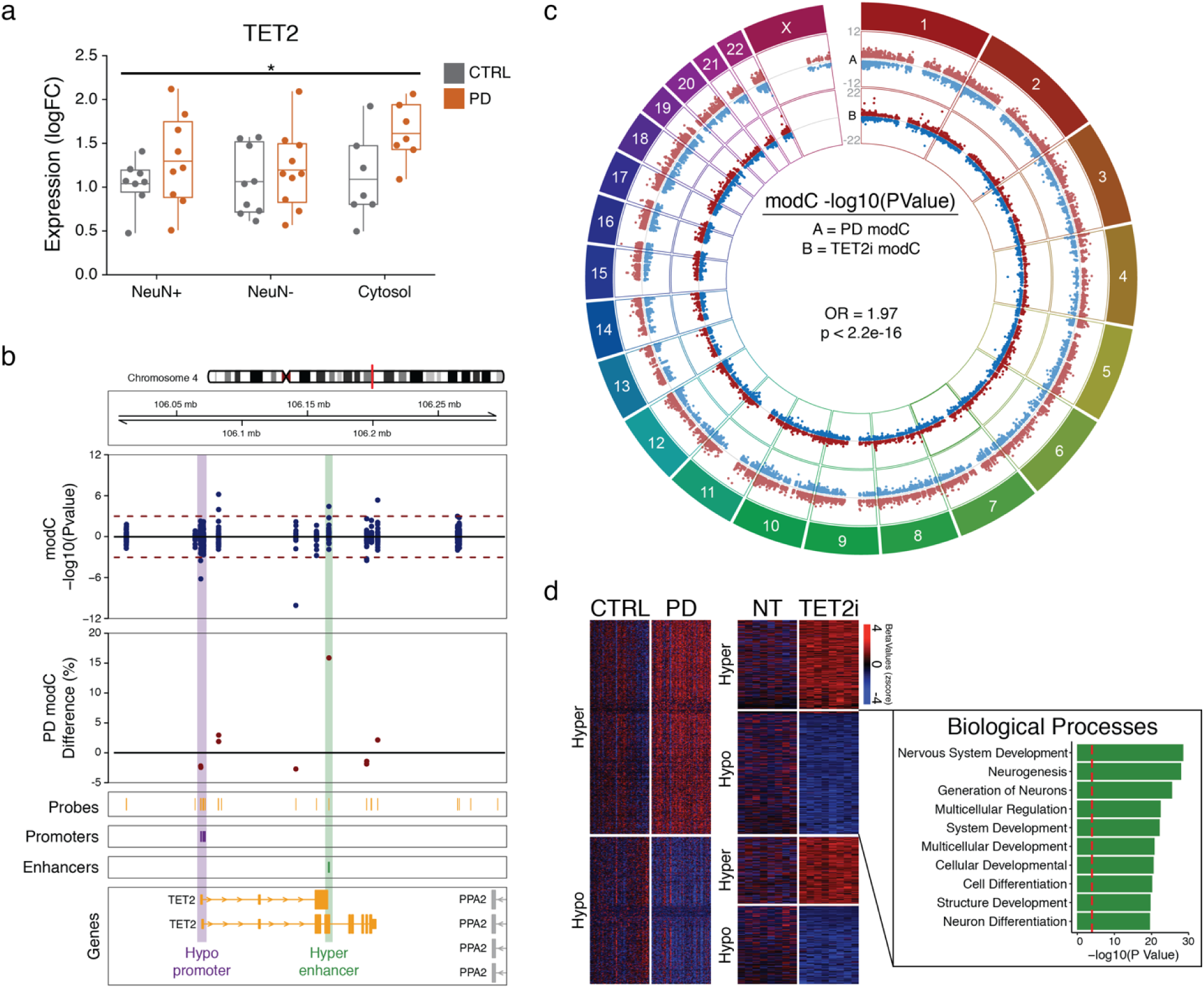
*TET2* transcript levels are upregulated in PD and depletion of *TET2 in vitro* induces cytosine modification changes that are inverse to those observed in PD neurons. (**a**) *TET2* transcript levels in PD neurons (NeuN+), glia (NeuN-), and bulk cytosolic space (cytosol) of PD patients (orange) and healthy controls (gray). \**p*<0.05 main effect of PD diagnosis determined by repeated-measures ANOVA. Boxplot center line is the mean, the lower and upper limits are the first and third quartiles (25th and 75th percentiles), and the whiskers are 1.5× the interquartile range. (**b**) DNA methylation analysis of the *TET2* genomic area. Differentially modified cytosines were identified in PD neurons using logistic regression, adjusting for age, sex, postmortem interval and neuronal subtypes (n=20 PD, 10 Controls, *q*<0.05). *TET2* promoter showing hypomethylation (purple) and enhancers (green) showing hypermethylation are highlighted. Promoters and enhancers in prefrontal cortex from NIH Roadmap Epigenomics Project (n=1)^28^. (**c**) The contribution of *TET2* to cytosine modification differences in PD neurons. Cytosine modification changes at enhancers in response to *TET2* inhibition by siRNA (TET2i), relative to non-target control, were profiled an *in vitro* model of neuronal cells, the SH-SY5Y neuroblastoma cell line. Differentially modified cytosines induced by *TET2* inactivation were identified using logistic regression (n=8 TET2i, 8 NT:non-targeting). Odds ratio (OR) and *p*<10^-15^ indicate that cytosine modification differences induced by *TET2* siRNA (A, light red: hypermethylated sites, light blue: hypomethylated sites) significantly overlap differentially methylated cytosines in enhancers in PD neurons (B, red: hypermethylated sites, blue: hypomethylated sites), as determined by Fisher’s exact test. (**d**) Heatmap and biological pathways for cytosine sites hypermethylated in PD neurons, and conversely, hypomethylated in response to *TET2* siRNA. Cytosine modification status for sites hypermethylated or hypomethylated in PD neurons (left panels; 1,799 cytosines at *q*<0.05 by logistic regression from Fig. 1a) were examined in the *TET2* siRNA SH-SY5Y cells (right panel). Biological pathways affected by genes with hypermethylated enhancers in PD neurons and hypomethylated enhancers in *TET2* inactivated SH-SY5Y cells (-log10 (PValue)). Pathway analysis by MetaCore with red line *q*<0.05, as determined by hypergeometric test.

We next explored the capacity for *TET2* to induce epigenetic changes at cytosine sites associated with PD in neurons. *TET2* mRNA levels were depleted by siRNA in an *in vitro* neuronal cell model, SH-SY5Y cells (1.41-fold loss in *TET2* mRNA in cells treated with siRNA cocktail, relative to non-targeting control; *p*<0.0001; unpaired equal variance t-test, Supplementary Fig. 6a). The effects of decreased *TET2* mRNA levels were profiled using the human brain enhancer bisulfite padlock probe library, which examined 1,031,160 modified cytosine sites. Overall, *TET2* depletion resulted in a significant decrease in cytosine modifications in the neuronal cell model (average 10.1% hypomethylation; OR=1.52, *p*<10^-15^, Fisher’s exact test; Supplementary Fig. 6b). When the differentially modified enhancers in PD neurons were examined, there was again a significant hypomethylation in *TET2* knockdown SH-SY5Y cells (74.3% of enhancer cytosine sites were hypomethylated; OR=1.97, *p*<10^-15^, Fisher’s exact test; **Fig. 5c**; Supplementary Data 13), which suggests that reduced *TET2* expression has the reciprocal effect to that observed in PD. In accordance, cytosine sites exhibiting a DNA modification gain in PD neurons (**Fig. 1a**) were also more likely to have a significant loss in DNA modification from *TET2* depletion in SH-SY5Y cells (74.3% of *q*<0.05 cytosines; *p*<10^-15^, Fisher’s exact test; **Fig. 5d**; Supplementary Data 14). Pathway analysis revealed that these sites affected neurogenesis and neurodevelopment (*q*<0.05, hypergeometric distribution; **Fig. 5d**). Together, these findings support that in PD neurons there is an upregulation of *TET2,* which leads to aberrant enhancer activation affecting neuronal differentiation pathways; a process that is reversed by *TET2* inhibition.

### Tet2 inactivation protects against inflammation-induced dopaminergic neuronal loss

We have observed that epigenetic activation of *TET2* is associated with PD. Therefore, we sought to determine whether loss of *Tet2* in mice was capable of limiting the development of neuropathology related to PD. Considering the abundance of proinflammatory transcriptional changes we observed in the PD brain, we examined the effects of *Tet2* inactivation in combination with an established inflammation model of PD; the lipopolysaccharide (LPS) challenge^46–49^. In this PD model, a single systemic dose of LPS generates a transient peripheral immune reaction that eventually leads to the progressive degeneration of nigral dopaminergic neurons^47^. The initial peripheral immune response to LPS lasts less than a week, but proinflammatory cytokines upregulated in the brain engage inflammation responses that gradually results in nigral neurodegeneration (apparent at ≥10 months post-injection)^47^. For this study, we administered one injection of LPS (5mg/kg, i.p.) or saline to 2-month old adult *Tet2* knock-out and wild-type mice, and then aged the mice to 11 months post-injection (n=7-14 mice per group; **Fig. 6a**). In the aged mice, we evaluated nigral dopaminergic neuronal loss, microglial activation, α-synuclein pathology, as well as motor behaviors (**Fig. 6a**).

**Fig. 6.**
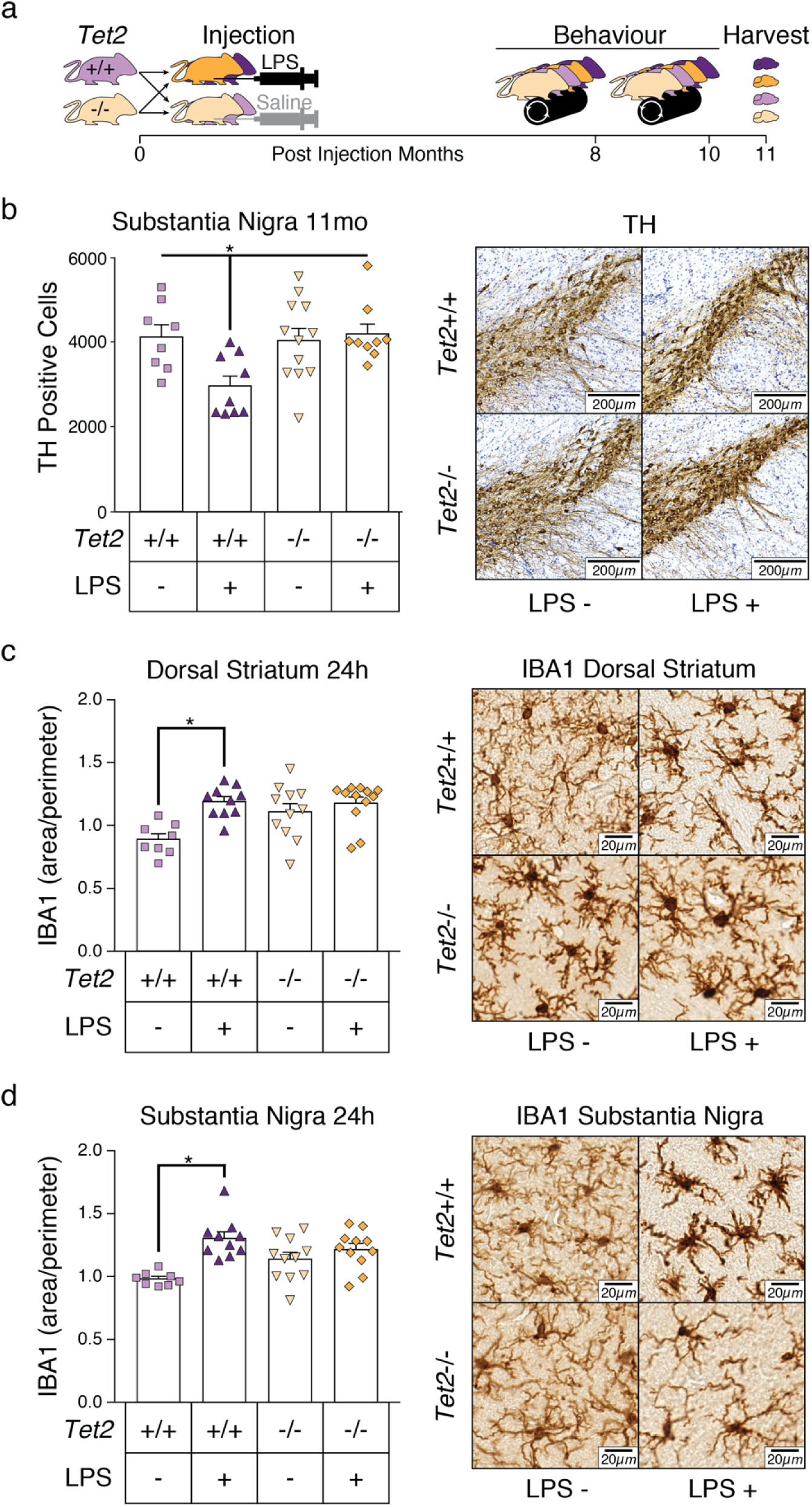
*Tet2* inactivation protects against nigral dopaminergic neuronal loss induced by previous exposure to systemic inflammation. (**a**) Schema of experimental design showing wild-type (*Tet2*+/+: purple) and *Tet2* knock-out (*Tet2*-/-: yellow) mice injected once with 5mg/kg LPS or saline at 2 months of age, followed by brain tissue analysis 11 months post-injection. (**b**) Immunohistochemistry and stereological analysis of TH-positive dopaminergic neuron counts in the substantia nigra of wild-type (+/+: orange) or *Tet2* knock-out (-/-: purple) mice injected with LPS or saline (n= 8 +/+ saline, 9 +/+ LPS, 12 -/- saline, and 9 -/- LPS). Representative images of substantia nigra dopaminergic neurons (TH-positive). Scale refers to 200 microns. (**c, d**) Microglial activation in the brain of *Tet2* knock-out and wild-type mice 24 h after exposure to LPS-mediated inflammation. Immunohistochemistry image analysis of microglial activation in the (**c**) dorsal striatum (n= 8 +/+ saline, 10 +/+ LPS, 11 -/- saline, and 12 -/- LPS) and the (**d**) substantia nigra (n= 8 +/+ saline, 10 +/+ LPS, 11 -/- saline, and 11 -/- LPS). Representative images of dorsal striatum and substantia nigra microglia (IBA1-positive). \**p*<0.05 by post hoc Tukey HSD test with error bars representing s.e.m.

We found that *Tet2* inactivation protected nigral dopaminergic neurons against LPS-induced inflammation. There was a 28.5% loss in TH-positive dopaminergic neurons in the substantia nigra of wild-type mice 11-months post-LPS injection, relative to saline-treated wild-type mice (genotype × treatment interaction: F(1,34)=5.85, *p*<0.05, two-way ANOVA; **Fig. 6b**). In contrast, *Tet2* knock-out mice with prior LPS exposure retained normal numbers of nigral dopaminergic neurons (**Fig. 6b**). The number of dopamine neurons in *Tet2* knock-out mice previously exposed to LPS were comparable to those of saline-treated wild-type and *Tet2* knock-out mice (**Fig. 6b**), indicative of a robust protective effect.

We also examined microglial morphology in the substantia nigra and striatum 24 h and 11 months post-injection. At 24 h post-injection, the microglia in the substantia nigra and dorsal striatum were significantly activated in wild-type mice treated with LPS, whereas *Tet2* knock-out mice treated with LPS had no significant change in microglial state (genotype × treatment interaction in substantia nigra: F(1,36)=6.96, *p*<0.05; in dorsal striatum: F(1,37)=4.95, *p*<0.05, two-way ANOVA; **Fig. 6c and 6d**). At 11 months post-injection, LPS treatment no longer had an effect on microglial morphology in wild-type and *Tet2* knock-out mice (Supplementary Fig. 7). There were also no significant differences in proteinase K-resistant α-synuclein pathology in any mouse group (Supplementary Fig. 7). In behavioral measures, prior LPS treatment induced a decrease in motor learning in wild-type mice, which was reversed by *Tet2* inactivation (*p*<0.05, repeated measures ANOVA; Supplementary Fig. 8). No other overt motor differences were observed (Supplementary Fig. 8).

To understand the biological mechanisms that enable the protective effects of *Tet2* inactivation against nigral neurodegeneration, we profiled transcriptional responses to LPS-induced inflammation in *Tet2* knock-out and wild-type mice 24 h post-injection. Transcriptomic analysis was performed using RNA-sequencing of the midbrain of wild-type and *Tet2* knock-out mice injected with either saline or LPS (n=4 wild-type saline, n=4 wild-type LPS, n=3 *Tet2* knock-out saline, n=3 *Tet2* knock-out LPS). There were 1,616 differentially expressed genes in wild-type mice 24 h after LPS treatment, while *Tet2* knock-out mice treated with LPS had 1,860 differentially expressed genes, relative to saline treated mice (LPS compared to saline treatment within each genotype; *q*<0.05, generalized linear regression with contrasts fit; Supplementary Data 15). Pathway analysis revealed that though immune cell activation and inflammation occurred in both wild-type and *Tet2* knock-out mice in response to LPS, the number of pathways affecting the immune system was severely attenuated in the *Tet2* knock-out mice (immune system pathways in wild-type mice 343, in *Tet2* knock-out mice 256; *q*<0.01, GSEA; **Fig. 7**; Supplementary Data 16). Moreover, wild-type mice treated with LPS exhibited alterations in tissue development, activation of pro-apoptotic signaling cascades, and mitochondrial dysfunctions (*q*<0.01, GSEA), which were not observed in the *Tet2* knock-out mice (**Fig. 7**). Hence, *Tet2* inactivation substantially attenuates the reactivity of the brain to proinflammatory triggers of PD neuropathology.

**Fig. 7.**
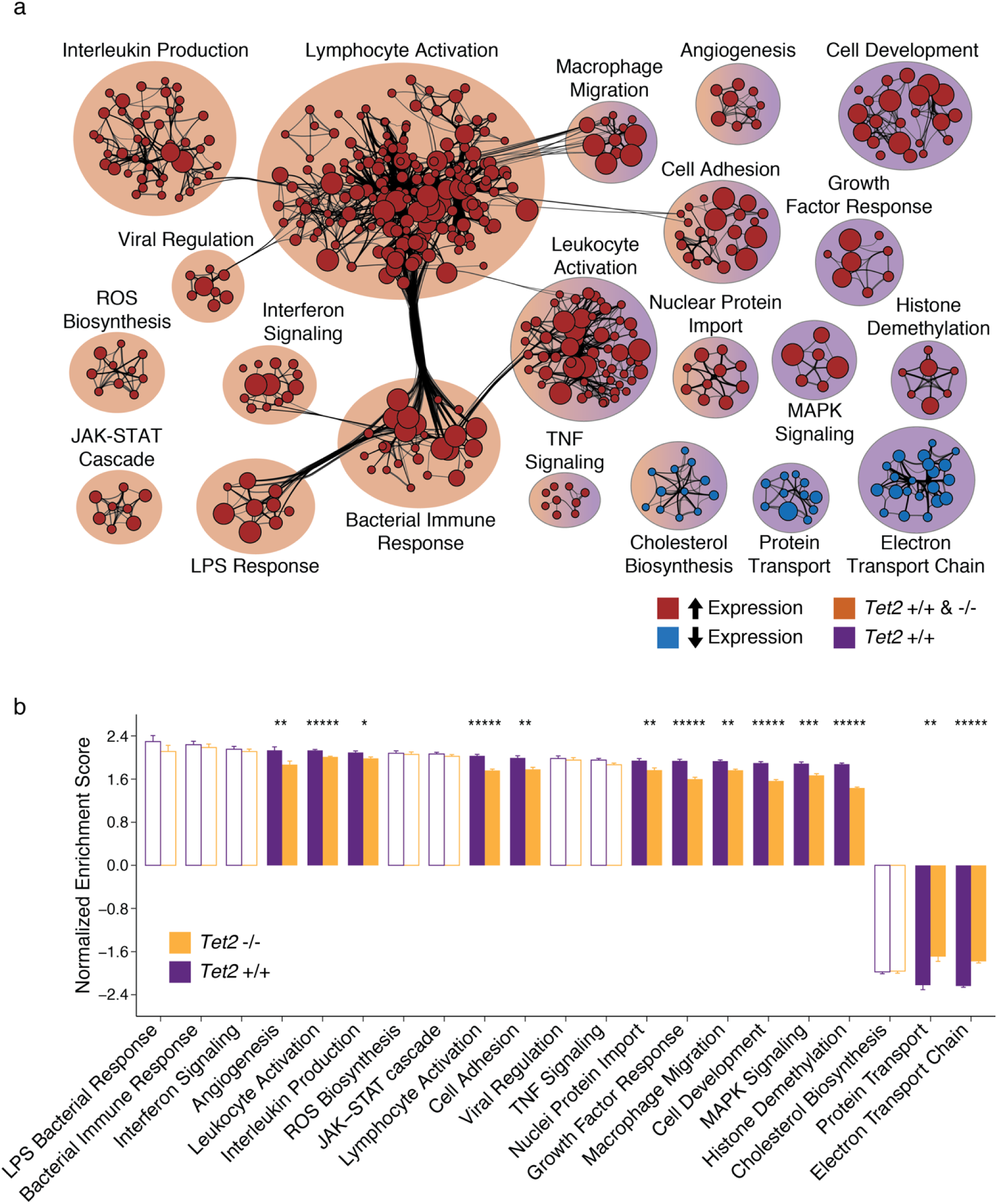
*Tet2* loss suppresses immune signaling pathways induced by proinflammatory stimulus. Differentially expressed genes induced 24 h post-LPS injection in wild-type and *Tet2* knock-out mice were identified using a generalized linear model (n=4 *Tet2*+/+ LPS, 4 *Tet2*+/+ saline, 4 *Tet2*-/-LPS, 4 *Tet2*-/- saline, *q*<0.05). (**a**) Pathway analysis of transcriptional changes induced by LPS in both the wild-type and *Tet2* knock-out mice (*Tet2* +/+ & -/-: orange) or in only the wildtype mice (*Tet2*+/+: purple). *q*<0.01 pathways, as determined by GSEA preranked. No pathways occurred in only the *Tet2* knock-out mice. Transcriptionally activated pathways in red and silenced pathways in blue. (**b**) Differential responses of *Tet2* knock-out mice relative to wild-type mice to LPS treatment. Normalized enrichment score of all subpathways identified to be altered by LPS injection in wild-type (purple) and *Tet2* knock-out (orange) mice. \**q*<0.05, \*\**q*<0.01, \*\*\*\**q*<10^-4^ by unequal variance T-test corrected for multiple testing. Error bars indicate s.e.m.

## Discussion

Our multiomics approach integrating epigenetics and transcriptomics in combination with analysis of *in vitro* and *in vivo* model systems provides new understanding of the molecular mechanisms involved in PD, and discovered a potential target for PD therapies. In an unbiased, genome-wide analysis of enhancers in PD neurons we identified a widespread increase in cytosine modifications, which involved a prominent contribution of hydroxymethylation. TET enzymes control levels of hydroxymethylation^23^, and we observed an epigenetic dysregulation of *TET2*, which was replicated in two separate epigenetic analyses of PD neurons. Moreover, we found a gain in *TET2* transcript levels in PD, and *in vitro TET2* manipulation in a neuronal cell model supported a contribution of *TET2* to the epigenetic abnormalities observed in PD neurons. Remarkably, suppression of *TET2* prevented nigral neurodegeneration induced by prior inflammation. This neuroprotective effect was found to be likely due a diminished inflammatory response in the brain. Overall, reduction in *TET2* levels was found to be neuroprotective and may be a novel avenue for therapeutic intervention in PD.

Recent genetic studies in neurons indicate that changes in enhancer activity may play an important role in many neurodegenerative diseases including PD^30, 50^. We showed that in PD patient neurons there is an increase in DNA modifications at enhancer CpG and CpH sites, which are both associated with a potent regulation of gene transcript levels^51^. Notably, our epigenetic analyses do not delineate epigenetic changes causally involved in PD from those that are a consequence of the disease processes. However, we find prominent increases in cytosine modifications prior to the arrival of pathology and at known PD risk genes, supporting that these epigenetic changes occur early in PD and may contribute to disease pathogenesis.

To further understand the types of cytosine modifications altered in PD neurons, we profiled hydroxymethylation, finding that a considerable proportion of cytosine modification increases in PD neurons (73%) were related to a rise in hydroxymethylation. Hydroxymethylation has been found to encompass as much as 10-40% of modified cytosine sites in neurons of the prefrontal cortex^52, 53^. Though DNA methylation at CpG and CpH sites in enhancers is largely regarded as gene silencing, hydroxymethylation is associated with active transcription and increased enhancer function^24, 53–55^. Hydroxymethylation, which is enriched at enhancers, has been shown to promote neuronal differentiation and accumulate in the aging brain^18, 23, 56, 57^. In accordance, we observed an enrichment of neurodevelopmental pathways altered by epigenetic changes at enhancers in PD. Activation of neurogenesis in postmitotic neurons of PD patients by the elevation of 5-hydroxymethylcytosine levels would have neurotoxic consequences, as postmitotic neurons undergo apoptosis in response to cell cycle reactivation^58^. A role for hydroxymethylation in neuronal dysfunction is further supported by reports of global increases in 5-hydroxymethylcytosine in multiple neurodegenerative diseases^59–61^, including in the PD brain^62^.

Consistent with the abnormalities in hydroxymethylation in PD neurons, we found an epigenetic disruption of *TET2* in PD neurons with a concomitant upregulation of *TET2* transcript levels. TET2 is abundant in the brain and impacts both neurogenesis and inflammatory pathways. TET2 activity is centrally involved in neuronal differentiation through the regulation of 5-hydroxymethylcytosine levels at neuronal enhancers^18, 23, 24, 63^. This signifies that increased *TET2* levels may be responsible for the epigenetic reactivation of neurogenesis pathways in PD, which prompts the death of terminally differentiated neurons^58^. In support, we found that depletion of *TET2 in vitro* impacted cytosine sites relevant to PD and involved in neurodevelopment. Moreover, induced inflammation has been previously shown to increase both TET2 activity and subsequent hydroxymethylation^64–66^. Hence, elevated levels of TET2 and accompanying hydroxymethylation may be a consequence of increased inflammation in the PD brain. However, there is a probable bidirectional relationship between TET2 activation and inflammation since intrinsic TET2 activity promotes immune cell development. TET2 regulates both the adaptive and innate immune systems by recruiting transcription factors important in immune cell differentiation and lineage specificity^42, 67, 68^, enabling inflammatory responses. Thus, TET2 disruption could substantially contribute to the epigenetic and transcriptomic pathway changes that converge on aberrant neuronal differentiation and prominent inflammatory activation in the PD brain (**Fig. 8**).

**Fig. 8.**
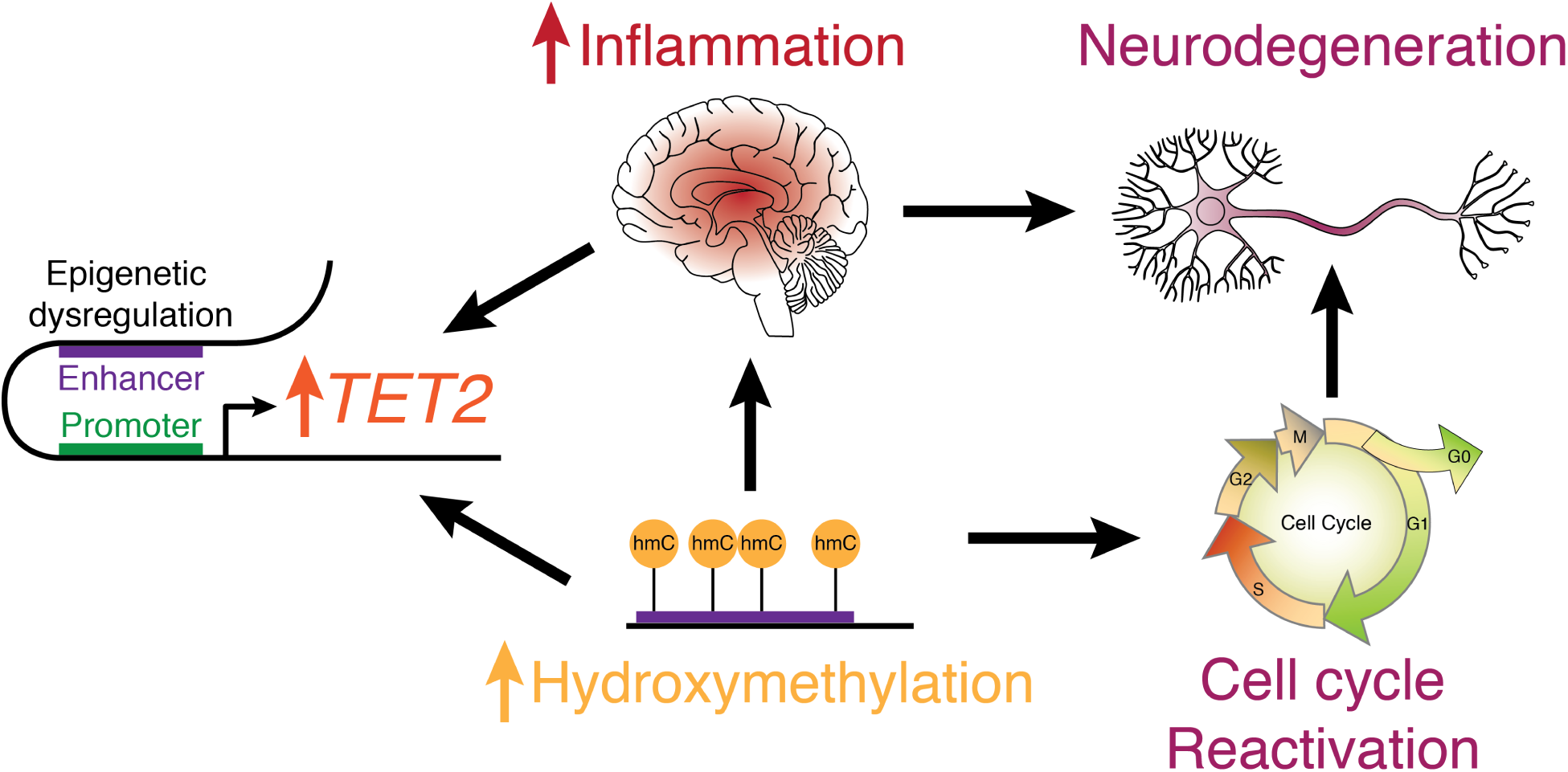
Proposed model displaying the contribution of *TET2* to the development and progression of PD. *TET2* is responsible for the conversion of DNA methylation to hydroxymethylation, and *TET2* inactivation results in cytosine modification changes relevant to those observed in PD neurons. In PD, there is epigenetic dysregulation of the *TET2* locus and an upregulation of *TET2* expression. This leads to increased hydroxymethylation, which has two main consequences. First, the hydroxymethylation increases susceptibility to immune activation, resulting in neuroinflammation and a positive feedback loop that further upregulates *TET2* activation. Second, hydroxymethylation reactivates genes involved in neurogenesis in post-mitotic neurons. Both of these aberrant processes may lead to the neurodegeneration observed in PD patients.

Our results show that inactivation of *TET2 in vivo* protected dopaminergic neurons from inflammation induced neurotoxicity. *TET2* inactivation also blunted transcriptional inflammation responses, and studies of hematological malignancies suggest that this could be related to a shift towards a more immature immune system^68–70^. Therefore, increased *TET2* expression in PD patients may result in hypervigilant immune activity along with a compromised neuronal system that leads to increased sensitivity to inflammation-induced neurotoxicity. Considering that neuroinflammation may further propel increased *TET2* expression, this suggests a potentially reinforced feed-forward loop contributing to the overall neuroinflammation and neurotoxicity observed in PD patients^42^ (**Fig. 8**). Currently, there are several ongoing clinical trials and a strong research interest in testing the therapeutic efficacy of preventing neuroinflammation in PD patients^71–73^. Our study suggests that compounds capable of inhibiting TET2 activity, like dimethyloxalylglycine (DMOG)^74, 75^, may be beneficial in PD treatment. In particular, DMOG has been shown to be neuroprotective after traumatic brain injury and hypoxia^76^. Together, our results suggest that reducing *TET2* in PD patients may be a new therapeutic target, which through the regulation of the epigenome could dampen an overactive immune system and protect against neurodegeneration.

## Methods

No statistical methods were used to predetermine sample size.

### Human Tissue Samples

Prefrontal cortex tissue of our sample cohort were obtained through the Parkinson’s UK Brain Bank, NIH NeuroBioBank, and Michigan Brain Bank. For each individual, we obtained information on demographics (age, sex), tissue quality (post-mortem interval, tissue quality/RIN score) and clinical variables (PD diagnosis, PD Braak stage, Alzheimer’s disease beta-amyloid pathology, and Alzheimer’s disease Braak stage and tau pathology). Patient data is provided in Supplementary Data 1. Our study included 105 individuals: 57 PD and 48 controls. Our linear regression models controlled for age, sex, tissue quality, and clinical variables, with each model checking variance inflation factors (VIF ≤ 5) to verify lack of multicollinearity. The study protocol involving use of postmortem brain tissue was approved by the ethics committee of the Van Andel Research Institute (IRB #15025).

### Neuronal Nuclei Isolation

Isolation of neuronal nuclei from prefrontal cortex was performed using a flow cytometry-based approach, similarly to previously described^77, 78^. Briefly, human brain tissue (∼250 mg) for each sample was minced in 2 mL PBSTA (0.3 M sucrose, 1X phosphate buffered saline (PBS), 0.1% Triton X-100) and homogenized in PreCellys CKMix tubes with a Minilys (3,000 rpm for 5 s, 5 min on ice, 3 times) (Bertin Instruments). Samples homogenates were filtered through Miracloth (EMD Millipore), rinsed with 2 mL of PBSTA and placed on a sucrose cushion (1.4 M sucrose). Nuclei were pelleted by centrifugation at 4000 × g for 30 min 4°C using a swinging bucket rotor and the pellet was incubated in 700 μl of 1X PBS on ice for 20 min. The nuclei were then gently resuspended and blocking mix (100 μl of 1X PBS with 0.5% BSA (Thermo Fisher Scientific), and 10% normal goat serum (Gibco) was added to each sample. Anti-NeuN antibody with Alex Fluor 488 (1:500; Abcam; ab190195) was added and samples were incubated 45 min at 4°C with gentle mixing. Immediately prior to flow cytometry sorting, nuclei were stained with 7-AAD (Thermo Fisher Scientific) and passed through a 30 μM filter (SystemX). Nuclei positive for 7-AAD and either NeuN+ (neuronal) or NeuN‒ (non-neuronal) were sorted using a MoFlo Astrios (Beckman Coulter) running Summit 6.3 by the Van Andel Research Institute Flow Cytometry Core and by a BD Influx (BD Biosciences) in the Jovinge lab. Approximately 1 million NeuN+ nuclei were sorted for each sample with an average purity of 97.38% ± 0.33%. After flow cytometry sorting, NeuN+ nuclei were placed on ice, resuspended in a 0.3 M sucrose, 4.2 mM CaCl2 and 2.5 mM Mg(Ace)2 solution, and centrifuged at 1,786 × g for 15 min at 4°C. The resulting neuronal nuclei pellets were stored at -80°C prior to standard phenol-chloroform genomic DNA isolation.

### Fine-mapping of modified cytosines at enhancers with bisulfite padlock probes

DNA methylation at human brain enhancers were analyzed at single nucleotide resolution from prefrontal cortex neuronal nuclei using the bisulfite padlock probe technique^77, 79^. Human brain enhancers were identified using the EpiCompare tool^80^, which predicts tissue/cell specific enhancers and promoters from chromatin state data defined by the RoadMap Epigenomics Project ChromHMM tool (v1.14)^81^. All enhancers identified in the 18-state ChromHMM model were used to identify enriched brain-specific enhancers, using a Fisher’s exact test, against non-brain tissues and can be downloaded from the Tissue Specific Enhancers website (https://epigenome.wustl.edu/TSE/index.php). In addition to these brain-specific enhancers, we included all enhancers identified in the 18-state ChromHMM adult prefrontal cortex (E073), interior temporal lobe (E072), and substantia nigra (E074). Promoters, which at times can act as enhancers^29^, identified in the 18-state ChromHMM adult prefrontal cortex, interior temporal lobe, and substantia nigra, were also included.

Padlock probes were designed to target enhancer locations on both DNA strands of the GRCh37/hg19 bisulfite-converted genome. Padlock probes were generated using the ppDesigner software (v1.1)^82^. In total, there were 59,009 brain enhancer probes targeting the unique (non-repetitive) genome. Probe genomic locations and sequences are described in Supplementary Data 17. Probes were synthesized using a programmable microfluidic platform (Custom Array, Inc.) and prepared as previously described^77, 79^. Fine-mapping of DNA methylation at enhancers using bisulfite padlock probes was performed^77, 79^. For each sample, genomic DNA was bisulfite converted and purified using the EZ DNA Methylation-Lightning Kit (Zymo Research). The bisulfite-converted DNA (200 ng) was then hybridized to the padlock probes (1.5 ng). PfuTurbo Cx (Agilent Technologies) was used to extend across the target regions and circularization was completed using Ampligase (Epicentre). Non-circularized DNA was digested using an exonuclease cocktail and the remaining circularized DNA was amplified using a common linker sequence in the padlock probe. Libraries were PCR amplified, pooled in equimolar amounts, purified by QIAquick Gel Extraction kit (Qiagen) and quantified by qPCR (Kapa Biosystems) on a ViiA 7 Real-time PCR system (Applied Biosystems). Next-generation sequencing of the libraries was performed on an Illumina HiSeq 2500 machine on HiOutput mode across 3 flow cells (24 lanes), producing ∼30 million reads per sample, at the Epigenetics Lab at the Centre for Addiction and Mental Health in Toronto, Canada.

### Mapping modified cytosines at *TET2*

DNA methylation was profiled in-depth at *TET2* in prefrontal cortex neurons of 20 PD patients and 11 controls. Padlock probes were designed to target the entire *TET2* locus and surrounding area (± 300 kb) using the ppDesigner software^82^ with the GRCh37/hg19 bisulfite-converted genome. Probes synthesis, library preparation, and bisulfite sequencing was performed as described in the Fine-mapping of modified cytosines at enhancers with bisulfite padlock probes Methods Section.

### Enhancer DNA methylation analysis

A custom pipeline based on the Bismark tool^83^ was used to interrogate every cytosine site (CpG and CpH) at 33,249 enhancer regions covered by padlock probes. In data preprocessing, adapter sequences were removed using Trimmomatic (v0.32), and sequencing reads aligning to the phiX DNA spiked-in were removed. The remaining sequencing reads were then aligned to the target reference genome (GRCh37/hg19) and DNA methylation calls were made for cytosines with a read coverage of at least 30× using Bismark (v0.17.0). At each cytosine site, DNA methylation calls were determined as the percent fraction of reads that retained the reference “C” (not converted to “T” by the bisulfite treatment). Bisulfite conversion efficiency was calculated to be 99.14 ± 0.05% (averaged CpA methylation per sample). Sequencing and technical replicates confirmed that there was a high reproducibility in sample-level DNA methylation, using a correlation analysis (average R for: Sequencing replicates (0.978 ± 0.00104 s.e.m), Technical replicates (0.972 ± 0.0012 s.e.m) (Supplementary Fig. 2). Two replicate samples were removed due to poor inter-sample correlation (R < 0.9). DNA methylation calls at each site were then merged across matched replicate samples (n=28).

Cytosine sites with missing DNA methylation calls in more than 30% of samples or that lacked DNA methylation (0% DNA methylation status) across all samples were excluded from further analysis. Cytosine sites overlapping common SNPs, identified by the 1000 Genomes Project (phase 3 v5a 20130502 release for chr1∼chr22, v1b 20130502 for chrX; African, European and all populations)^84^, were also excluded from further analysis. After data preprocessing, 904,511 cytosine sites, including both CpG and CpH sites, had quality-controlled genome-wide DNA methylation calls for 105 individuals.

In our DNA methylation analysis of prefrontal cortex neuronal nuclei, we controlled for inter-sample variation in neuronal subtypes, glutamatergic and GABAergic (GABA) neurons. In the prefrontal cortex, excitatory glutamatergic neurons account for 70–85% of neurons, while the remaining 15–30% are inhibitory GABA neurons^85^. Glutamatergic and GABA neurons have 12 and 9 subtypes, respectively^35^. We verified that there was not a selective loss of any of the 21 specific neuronal subtypes. In our dataset, we averaged CpH methylation within gene bodies (±100 kb) and found 582 neuronal subtype-specific gene signatures. Cell-type deconvolution was performed using CIBERSORT^36^ (http://cibersort.stanford.edu; 100 permutations) with the 582 neuronal subtype-specific signatures. There were no significant differences identified between PD and control individuals in any of the 21 specific neuronal subtypes (Supplementary Fig. 3). In our subsequent analysis, we adjust for neuronal subtype proportion, which refers to the proportion of glutamatergic to GABA neuronal subtypes.

Our DNA methylation analysis identified CpG and CpH sites associated with PD in prefrontal cortex neuronal nuclei. The counts and coverage data for the 904,511 DNA methylation sites were inputted into methylKit (v1.8.1)^86^, and a logistic regression model controlling for age, sex, post-mortem interval, and neuronal subtype proportion, was used to identify differentially methylated sites in neurons of PD patients compared to controls. Multiple testing correction was performed using the Benjamini-Hochberg method. A Fisher’s exact test was used to determine enrichment of hypermethylation at the differential methylated sites in PD. Transcription factor (TF) analysis was performed using oPOSSUM (v3.0)^87^. To identify significantly overrepresented TF families in enhancer regions altered in PD, we performed the TF Binding Site (TFBS) Cluster Analysis in oPOSSUM, using the differentially methylated sites in PD (± 20 bp) against all examined DNA methylation sites (± 20 bp) as background.

### Gene annotation and enrichment analysis

Enhancers dramatically affect gene expression through chromatin interactions with gene promoters. To identify the gene targets of the epigenetically misregulated enhancers in PD neurons, we analyzed human prefrontal cortex Hi-C data^41^ (Illumina HiSeq 2000 paired-end raw sequence reads; n = 1 sample; 746 million reads; accession: GSM2322542). Hi-C analysis used Trimmomatic (v0.4.4) to remove adapter sequences, HiCUP (v0.5.9)^88^ to align reads to the human reference genome (GRCh37/hg19), HOMER^89^ to identify significant genomic interactions (40 kb resolution, *p* < 0.001 and z score > 1.0) and HiCSeg (v1.1)^90^ to identify topological associated domains (TADs). Our analysis of the Hi-C data identified 58,758 interactions in total, of which 951 interactions involved the epigenetically altered enhancers in PD neurons. Gene targets of the enhancers were defined as gene promoters (± 2 kb of a gene TSS) interacting with an enhancer location. The Hi-C analysis determined that all enhancer locations profiled in our study interacted with 5,746 genes (background), of which 1,469 genes interacted with at least one differential methylation site in PD. To further identify proximal gene-enhancer interactions, the GREAT software^91^ was used. In total, we had 9,957 genes associated to enhancers profiled in our study (background) of which 2,885 genes interacted with at least one differential methylation site in PD.

Pathway enrichment analysis for genes affected by differential methylation at enhancers in PD neurons was performed using MetaCore (https://clarivate.com/products/metacore/). Comparison of pathways altered by differential methylation and differential mRNA expression in PD were determined using GProfiler^92^ and Gene Set Enrichment Analysis (GSEA)^93^, respectively. Cytoscape (v3.1.7), with EnrichmentMap (v3.1.0) and AutoAnnotate (v1.3) modules^94^, was used to cluster and visualize the enriched pathways.

### Hydroxymethylation profiling by hMeDIP Sequencing

We examined hydroxymethylation in prefrontal cortex neurons of 21 PD patients and 23 controls (a subset of samples examined in the bisulfite padlock probe study above). Genomic DNA from human prefrontal neurons was sheared using a Covaris LE220 sonicator to produce fragments in a size range of 200–600 bp. hMeDIP was performed using the EpiQuik kit according to the manufacturer’s instructions (Epigentek). Briefly, a 5-hmC antibody or non-immune IgG (i.e. negative control) was coated on the bottom of a 96-well plate. Wells were washed and either 1 µg sheared DNA, 1 µg control DNA, or buffer only was added to appropriate wells. The plate was then covered and incubated for 90 min with orbital rotation at 100 rpm. Wells were then washed 5 times with washing buffer. Captured DNA was then eluted by incubation of each well with a proteinase K mixture at 60°C for 15 min followed by incubation at 95°C for 3 min. The solution containing eluted DNA was transferred to another 96-well plate. DNA in each well was then purified using AMPure XP beads (Beckman Coulter). Fold-enrichment of target 5-hmC DNA was determined to be 950-fold, as determined by qPCR of the 5-hmC and non-immune IgG pulldown of known hMeDIP target sequence.

Sequencing libraries for input control and hMeDIP samples were prepared by the Van Andel Genomics Core from 10 ng of input material and all available immunoprecipitated material using the KAPA Hyper Prep Kit (v5.16) (Kapa Biosystems). End repaired and A-tailed DNA fragments were ligated to IDT for Illumina TruSeq UD Indexed Adapters (Illumina), followed by PCR amplification. Quality and quantity of the finished libraries were assessed using a combination of Agilent DNA High Sensitivity chip (Agilent Technologies), and QuantiFluor dsDNA System (Promega Corp.). Sequencing (single-end 100 bp) was performed on an Illumina NovaSeq6000 sequencer producing ∼30 million reads per sample. Base calling was done by Illumina RTA3, and output was demultiplexed and converted to FASTQ format with Illumina Bcl2fastq2 v2.19.

Adapter sequence from raw sequencing reads were removed using TrimGalore (v0.4.4). Sequenced reads from hMeDIP immunoprecipitated and input control human brain nuclei were mapped to the human reference genome (GRCh37/hg19) with Bowtie2 (v2.3.1)^95^. A combination of Picard and Samtools (v1.9) were used to mark and remove PCR duplicates respectively. For each sample, Deeptools (v2.3.1)^96^ was used to apply a Signal Extraction Scaling (SES)^97^ factor normalized to the input (1 kb wide tiling windows) prior to peak calling. Broad peaks were called using MACS2 (2.1.2)^98^ for each normalized sample and a consensus peak list was then defined as contiguous regions covered by at least 50% of samples. Consensus peaks in the shortest 5% and longest 5% range were removed from further analysis. In total, we identified 273,519 hydroxymethylation consensus peaks (*q*<0.05). For each consensus peak, bedtools maps were used to extract SES normalized counts. To determine hydroxymethylation contribution to epigenetic changes at enhancers in PD neurons, hydroxymethylation counts for the differentially methylated cytosines in PD were extracted using bedtools coverage (v2.25.0) and TMM normalized. Counts less than one were marked as 0. Logarithmically transformed counts were passed into a generalized linear regression model with diagnosis, age, sex, postmortem interval, and neuronal subtype proportion as covariates. A multiple testing correction was applied using the Benjamini-Hochberg method. Fisher’s exact tests were used to determine the enrichment of hydroxymethylation and gain in hydroxymethylation.

### RNA Sequencing

RNA sequencing analysis was performed on human prefrontal cortex tissue (n = 24 PD, 12 age-matched healthy controls). Frozen tissue (20–30 mg) was homogenized in TRIzol (Invitrogen) using a Precellys Lysing Kit (Bertin Technologies) with the MiniLys (Bertin). Total RNA was isolated and treated with DNase I (Qiagen), followed by a cleanup using a RNeasy Mini Kit (Qiagen). The resulting RNA quantity was assessed by Nanodrop 8000 (Thermo Scientific) and quality was assessed with an Agilent RNA 6000 Nano Kit on a 2100 Bioanalyzer (Agilent Technologies). RNA library preparations from 500 ng of total RNA per sample was performed by the Van Andel Genomics Core. RNA libraries were prepared using the KAPA Stranded KAPA RNA HyperPrep Kit with RiboseErase (v1.16) (Kapa Biosystems), as per manufacturer’s instructions. In brief, RNA was sheared to 300–400 bp, cDNA was converted, cDNA fragments were ligated to Bioo Scientific NEXTflex Adapters (Bioo Scientific), and PCR amplified. RNA sequencing libraries were assessed for quality and quantity using Agilent DNA High Sensitivity chip (Agilent Technologies, Inc.) and the QuantiFluor dsDNA System (Promega Corp.), respectively. Kapa Illumina Library Quantification qPCR assays (Kapa Biosystems) were used to pool equal equimolar concentrations of the individually indexed libraries, prior to single-end 75 bp sequencing on an Illumina NextSeq 500 sequencer producing ∼40 million reads per sample.

In our RNA sequencing analysis pipeline, adapter sequences were removed using TrimGalore (v0.4.4) prior to genomic alignment. STAR (v2.3.5a)^99^ was used to align and count sequencing reads to either human (GRCh37/hg19) Gencode v19 primary assembly transcriptome. Gene counts matrix was imported into R (3.5.1) and low expressed genes (counts per million < 1 in all samples) were removed prior to trimmed mean of M-values normalization in edgeR (v3.24.3). Transcriptomic analysis to identify differentially expressed genes in PD relative to controls was performed using a generalized linear model within edgeR^100^, adjusting for age, sex, RIN, and sources of unknown variation. Sources of unknown variation was determined using stable control genes (*p* ≥ 0.5) in the RUVseq Bioconductor package (v1.16.1)^101^.

To identify transcript level changes significantly associated with both PD and enhancer DNA methylation status, a linear modeling approach was adopted using limma (v3.38.3). For each individual, the mRNA expression values were adjusted for age, sex, RIN, and sources of unknown variation (gene expression residuals). DNA methylation values were adjusted for age, sex, postmortem interval, and neuronal subtype proportion (DNA methylation residuals). An empirical bayes linear model^102^ comparing gene expression changes with DNA methylation residuals, adjusted for diagnosis, allowed for the identification of genes whose expression significantly correlated with the associated enhancer DNA methylation status. Furthermore, in the same linear model comparing gene expression changes with PD diagnosis adjusted for DNA methylation changes, we identified genes associated with PD diagnosis. Intersections of both significant lists identified genes whose expression is associated with both DNA methylation status and PD diagnosis.

### *TET2* transcript levels in brain nuclei

*TET2* mRNA levels in neuronal nuclei, non-neuronal (glial) nuclei, and total cytoplasm was analyzed in 10 PD patients and 10 controls. To collect RNA, the neuronal nuclei isolation protocol was performed with minor modifications. In brief, Superase-In RNase Inhibitor (Thermo Fisher Scientific) was added to the PBSTA buffer and blocking mix at a 1:200 dilution. Total cytoplasm was harvested from the supernatant after pelleting the nuclei (through the 1.4 M sucrose cushion). Neuronal nuclei and non-neuronal nuclei were isolated by flow cytometry. Nuclei and total cytosolic fractions were stored at -80°C prior to qPCR. Neuronal nuclei and non-neuronal nuclei pellets were resuspended in PBS (∼8000 nuclei/uL). qPCR was performed using the TaqMan Gene Expression Cells-to-CT kit (Ambion) according to manufacturer’s instructions (lysis step removed for the cytosolic fraction). Each sample was run in triplicate on an Applied Biosystems StepOne Plus qPCR system. ΔΔCT value for *TET2* levels were relative to RPLP0 housekeeping gene (Thermo Fisher Scientific). Primers sequences are listed in Supplementary Data 17.

### Enhancer methylation changes in neurons induced by *TET2* siRNA

We determined whether altered *TET2* mRNA levels induce, in neurons, differential methylation at enhancers relevant to PD, using RNA interference in the SH-SY5Y neuroblastoma cell line (ATCC). For this approach, ∼1 million SH-SY5Y cells (∼70 % confluent well of a 6-well plate), cultured in 10% FBS (Gibco) RPMI (Thermo Fisher Scientific), were transfected with 100 nM of either human *TET2* siGENOME SMARTpool siRNA or a non-targeting siGENOME Control#2 siRNA (Dharmacon), using X-tremeGENE siRNA Transfection Reagent (Sigma-Aldrich) according the manufacturer’s instructions. After 72 h, the cells were pelleted and stored at -80°C prior to genomic DNA isolation, or homogenized in Trizol (Thermo Fisher Scientific) and stored at -80°C prior to RNA isolation. Genomic DNA was extracted by proteinase K digestion and standard phenol:chloroform extraction methods. Total RNA was isolated using Trizol (Thermo Fisher Scientific), followed by an on-column DNase treatment and RNA clean-up using the RNeasy kit (Qiagen), according to manufacturer’s instructions. cDNA was synthesized from 2 ug of total RNA using SuperScript IV One-Step RT-PCR System (Invitrogen). The efficacy of *TET2* knock-down was determined by qPCR using Power SYBR Green Master mix (Applied Biosystems) on a Quantstudio 3 Real Time PCR system (Thermo Fisher Scientific). Each sample was run in triplicate. *TET2* mRNA expression was determined using the ΔΔCT, with three housekeeping genes (*B2M*, *HPRT*, *RPL13*). Primers sequences are listed in Supplementary Data 17.

To profile DNA methylation at enhancers in SH-SY5Y cells exposed to *TET2* siRNA or non-targeting siRNA control, we used the same bisulfite padlock probes and library preparation approach described above. Sequencing was performed on Illumina HiSeq 2500 machine on HiOutput mode at the Epigenetics Lab at the Centre for Addiction and Mental Health in Toronto, Canada. Sequencing data preprocessing and quality control steps were performed as described above. All samples and replicates had a high inter-sample correlation (R > 0.95). DNA methylation calls at each site were merged across matched replicate samples. Cytosine sites with missing DNA methylation calls in more than 30% of samples or that lacked DNA methylation (0% DNA methylation status) across all samples were excluded from further analysis. After data preprocessing, 1,031,160 cytosine sites, including both CpG and CpH sites, had quality-controlled genome-wide DNA methylation calls for 4 *TET2* siRNA samples and 4 non-targeting siRNA control samples. The counts and coverage data for the 1,031,160 DNA methylation sites were inputted into methylKit^86^, and a logistic regression model was used to identify DNA methylation changes induced by depletion of *TET2* expression. Multiple testing correction was applied using the Benjamini-Hochberg method. Fisher’s exact test was used to determine the enrichment of differentially methylation sites induced by *TET2* knock-down in the previously identified differentially methylated sites in PD.

### Immunohistochemistry in *Tet2* knock-out mice

*Tet2* knock-out mice were obtained from The Jackson Laboratory (strain 023359). Heterozygote intercrosses were used to generate *Tet2* knock-out (*Tet2*-/-) and wild-type (*Tet2*+/+) mice, and genotyping was performed as previously reported^70^. All mice were maintained in a pathogen-free barrier facility and provided with food (LabDiet 5021) and water for *ad libitum* consumption. The vivarium was maintained under controlled temperature (21°C±1°C) and humidity (50-60%), with a 12-h diurnal cycle (lights on: 0700-1900). Animals were treated in strict accordance with the NIH Guide for the Care and Use of Laboratory Animals, with all animal experiments being approved by the Van Andel Institute Animal Care and Use Committee (AUP # PIL-17-10-010). Approximately equal numbers of male and female mice were used, and no sex differences were detected (in immunohistochemistry or behavioral testing). No animals were excluded from the study, and sample sizes were comparable to other studies of PD mouse models^103, 104^. At two months of age mice received an intraperitoneal injection of saline (PBS) or lipopolysaccharide (LPS; 5 mg/kg). Animals remained in home cages until tissue harvest or behavioral procedures were conducted. For the acute inflammation study, brain tissues were harvested exactly 24 h post-injection. For the chronic inflammation study, behavioral testing was performed at 8 and 10 month post-injection, and brain tissue harvest was at 11 months post-injection. Immunohistochemistry and behavioral studies in mice were conducted by an experimenter blind to genotype and treatment.

Mice were perfused with PBS and brain tissue for immunohistochemical analysis was harvested and fixed with 4% paraformaldehyde (pH 7.4) for 24 hours. After fixation, brains were cryoprotected in 30% sucrose for at least 48 hours and dissected into 40-μm-thick coronal sections. Prior to immunostaining for α-synuclein aggregates, brain sections were treated with 10 μg/ml proteinase-K (Thermo Fisher Scientific) in TE buffer (50 mM Tris-HCl pH 8.0, 1 mM EDTA, 0.5% Triton X-100) for 10 min and washed 3 times for 10 min in PBS Buffer (10 mM sodium phosphate pH 7.2, 0.1% Triton X-100). Immunostaining of brain tissue sections was performed by incubation with primary antibody (1:500 mouse α-synuclein, Biosciences; 1:500 rabbit IBA1, Wako Pure Chemical Corporation; 1:1500 mouse TH, Millipore), followed by biotinylated secondary antibodies (Goat Anti-Rabbit, Vectorlab; Goat Anti-Mouse, SourthernBioTech), ABC reagent (Vector Labs), and DAB (Vector Labs) as previously described^104^. For the microglia analysis, images of mounted slides were captured using a ScanScope XT slide scanner (Aperio) at a resolution of 0.24 µm per pixel. Microglia in the dorsal striatum (DS, 6 sections per mice) and substantia nigra pars compacta (SNpc, 8 sections per mice) were quantified using a random forest machine learning pixel classifier using ilastik (v1.3.2), trained on representative images of *Tet2* wild-type mice treated with either PBS or LPS for 24 h^105^. This trained model was used to unbiasedly predict all pixels representing microglia staining. A custom CellProfiler (v3.1.5) pipeline was used to quantify pixels representing microglia (90% confidence score)^106^. Microglia activation was calculated as mean area/perimeter per 1000 µm^2^. Proteinase-K resistant α-synuclein aggregates were also quantified using a custom CellProfiler pipeline per 1000 µm^2^. Stereology analysis of TH positive dopaminergic neurons in the substantia nigra was performed using the optical fractionator probe of StereoInvestigator software (MicroBrightField Biosciences), as previously described^104^.

### Behavioral procedures

Behavioral procedures were conducted 8 months and 10 months following treatment with LPS or saline. Prior to all behavioral procedures, animals were acclimated to the testing room for a minimum of 60 min. Open-field, catwalk, and rotarod, in this order, were conducted on successive days. Behavioral testing was conducted exclusively during the animal’s light cycle. The behavioral equipment was cleaned with 70% ethanol between each testing session.

*Open Field:* Gross motor activity of animals was assessed in the open field using the Any-maze Behavioral Tracking system (San Diego Instruments). At the start of each session mice were placed in the center of a 50 cm X 50 cm plexiglass chamber and their movement was monitored continuously for 1 h. Following the session, locomotor activity was calculated automatically from the recorded video using the Any-maze Behavioral Tracking software (San Diego Instruments).

*CatWalk:* Gait of the animals was assessed using the CatWalk XT system (Noldus). Prior to testing, mice were briefly acclimated for 10 min and trained (∼10 min) to walk across the CatWalk. During testing, mice were placed on the CatWalk and allowed to crossover to the opposing end, which contained a small housing box with block diet. Each test consisted of 3 consecutive trials. Gait analysis was then performed using the automated gait analysis system (Noldus Information Technology).

*Rotarod:* Rotarod performance was assessed on a computerized rotarod system (San Diego Instruments). To allow for acclimation, mice were initially placed on the stationary horizontal rod for 10 min. Learned rotarod motor skills was assessed at 8 months post-LPS or saline treatment. For this, mice were placed on the rod and rotation began at a rate of 4 rpm with an acceleration rate of 4 rpm/min for 5 min. Mice were tested once a day for 3 successive days, with a 24 h interval between each test. In addition, at the 8- and 10-month interval after LPS or saline treatment, each animal underwent a rotarod motor behavior test, consisting of 4 rpm constant for 60 s followed by an acceleration rate of 4 rpm/min for 4 min. This test was conducted over consecutive 3 days with each mouse undergoing 3 sessions per day. Between each session, mice were placed in their home cage for a minimum of 1 h. For all tests, time from the start of the session to the time the animal fell (latency) was recorded automatically by light beam break. The animals mean latency for each day was used for subsequent analysis.

### Transcriptomic analysis of LPS-treated *Tet2* knock-out mice

Transcriptomic analysis of the mouse midbrain 24 h after LPS treatment (5 mg/kg i.p.) was performed to determine the effects of inflammation on the midbrain of mice with *Tet2* inactivation (n = 4 *Tet2*+/+ mice treated with LPS, 4 *Tet2*+/+ mice treated with saline, 3 *Tet2-/-* mice treated with LPS, 3 *Tet2-/-* mice treated with saline). RNA isolation, sequencing, and data preprocessing with alignment to the mouse (GRCm38/mm10) Gencode vM20 primary assembly transcriptome was performed as in the RNA-sequencing Methods section, with minor modification. RNA libraries were prepared using the KAPA Stranded mRNA-seq library preparation kit (v5.17) (Kapa Biosystems), which included RNA shearing to 200–300 bp prior to cDNA synthesis. Single-end 75 bp sequencing on an Illumina NextSeq 500 sequencer produced ∼30 million reads per sample. For transcriptomic analysis in *Tet2* knock-out and wild-type mice treated with saline or LPS, we used a generalized linear model with contrast fits, adjusting for sources of unknown variation. Sources of unknown variation was determined using stable control genes (*p* ≥ 0.5) in the RUVseq Bioconductor package (v1.16.1)^101^.

### qPCR analysis of LPS-treated *Tet2* knock-out mice

qPCR analysis of inflammatory markers in the mouse midbrain 11 months after LPS treatment (5 mg/kg i.p.) was performed (n = 8 *Tet2*+/+ mice treated with LPS, 6 *Tet2*+/+ mice treated with saline, 10 *Tet2-/-* mice treated with LPS, 5 *Tet2-/-* mice treated with saline). Total RNA was isolated as described in the RNA-sequencing Methods section and converted to cDNA using the High Capacity cDNA Reverse Transcription Kit (Thermo Fisher Scientific). qPCR was performed using TaqMan Gene Expression Master Mix (Applied Biosystems) and Taqman Gene Expression Assays (Thermo Fisher Scientific) for the following genes: TNF-α, RELA, IL-6, NFKB2, and IL-1β (primers in Supplementary Data 17). Sample were run in triplicate on an Applied Biosystems StepOne Plus qPCR system. ΔΔCT value for target genes relative housekeeping genes (HPRT and GAPDH; Thermo Fisher Scientific).

## Acknowledgments

We thank the Van Andel Research Institute Flow Cytometry, Genomics and Bioinformatics, Pathology, and Vivarium Cores. We also thank the CAMH Sequencing Facility. We thank the Parkinson’s UK Brain Bank, the NIH NeuroBioBank, and the Michigan Brain Bank for the brain tissue provided. V.L. is supported by grants from the Department of Defense (W81XWH1810512), the National Institute of Neurological Disorders and Stroke (1R21NS112614-01), a Gibby & Friends vs. Parky Award, and a Van Andel Research Institute Innovation Award. SJ and MW are supported by the Richard and Helen DeVos Foundation.

## Data Availability

All sequencing data generated in this study are available from the NCBI Gene Expression Omnibus (GEO) database under the accession number GSE136010. RNA-seq data used in this study is available under the GEO accession number GSE135037. Custom code for DNA methylation and RNA-seq analysis is available at https://github.com/LeeLMarshall/Epigenomic-analysis-of-Parkinson-s-disease-neurons-identifies-Tet2-loss-as-neuroprotective.

## Author contributions

LM contributed to experimental design, computational analyses, immunohistochemistry experiments, and cell culture study. BK performed neuronal nuclei isolations, the bisulfite padlock probe sequencing of human patient samples and was involved in the immunohistochemistry and behavioral analyses in mice. PL was involved in the Hi-C analysis and experimental design of computational approaches. EE performed the *TET2* mRNA analysis in patient samples, RNA isolation of mouse samples, and contributed to the immunohistochemistry experiments. WC performed the bisulfite padlock probe sequencing of the cell culture study. KL contributed to the immunohistochemistry analysis. MW and SJ were involved in the flow sorting of neuronal nuclei. JG contributed to the hMEDIP analysis. GC contributed to experimental design. VL was involved with study design and overseeing the experiments. The manuscript was written by VL and LM and commented on by all authors.

## Competing interests

The authors declare no competing interests.

